# Feline leukemia virus (FeLV) endogenous and exogenous recombination events result in multiple FeLV-B subtypes during natural infection

**DOI:** 10.1101/2021.03.01.433497

**Authors:** Katelyn Erbeck, Roderick B. Gagne, Simona Kraberger, Elliott S. Chiu, Melody Roelke Parker, Sue VandeWoude

## Abstract

Feline leukemia virus (FeLV) is associated with a range of clinical signs in felid species. The primary hosts of FeLV are domestic cats of the Felis genus that also harbor endogenous FeLV (enFeLV) elements stably integrated in their genomes. EnFeLV elements display 86% nucleotide identity to exogenous, horizontally transmitted FeLV (FeLV-A). Variation between enFeLV and FeLV-A is primarily in the long terminal repeat (LTR) and env regions, which potentiates generation of FeLV-B recombinant subtypes during natural infection with enhanced virulence. The aim of this study was to examine exogenous FeLV (exFeLV) and enFeLV recombination events in a natural FeLV epizootic. We previously described that of 32 individuals in a closed colony with productive FeLV-A infection, 22 had detectable circulating FeLV-B. We cloned and sequenced the env gene of FeLV-B, FeLV-A, and enFeLV spanning known recombination breakpoints, examining between 1-13 clones per individual to assess sequence diversity and recombination sites. We documented multiple recombination breakpoints resulting in the production of unique FeLV-B genotypes. At least half of the cats harbored more than one FeLV-B variant, and almost all animals had variants similar to those recovered from at least one other individual in the colony. This analysis reveals that FeLV-B is predominantly generated de novo within each host, though horizontal transmission may be inferred based upon FeLV-B sequence identities between individuals. This work represents a comprehensive analysis of endogenous-exogenous retroviral interactions with important insights into host-viral interactions that underlie disease pathogenesis in a natural setting.

**Importance:** Feline leukemia virus (FeLV) is a felid retrovirus associated with a variety of disease outcomes. Exogenous FeLV-A is the most common horizontally transmitted virus subgroup. Domestic cats (*Felis catus*) harbor endogenous copies of FeLV (enFeLV) in their genomes. Recombination between FeLV-A and enFeLV may result in emergence of largely replication-defective, but highly virulent recombinant strains. FeLV-B, the most common recombinant form, results when enFeLV *env* recombines with FeLV-A during FeLV replication. This study evaluated endogenous-exogenous recombination outcomes in a naturally-infected closed colony of domestic cats to determine recombination sites and FeLV-B genotypic heterogeneity associated with enhanced disease virulence. While FeLV-A and enFeLV genotypes were highly conserved, a large number of unique FeLV-B variants were identified in association with predicted recombination hotspots. The findings provide insight into recombination events between viral and host genomes that result in new, and potentially more pathogenic, viral strains.

## Introduction

Endogenous retroviruses influence pathogenesis of exogenous viral infections by a wide variety of mechanisms (1, 2). An endogenous virus forms when proviral elements integrated into host germline cells and become a heritable component of the host genome. Over time, these endogenous viruses often acquire mutations that render them incapable of generating infectious virus (3-8). A prime example of this is feline leukemia virus (FeLV), a gammaretrovirus that infects domestic and wild felid species and is known to cause a range of diseases including lymphadenopathy, anemia, bone marrow suppression, immune suppression, lymphoma, leukemia, and ultimately, death (9, 10). FeLV has the highest case fatality rate in domestic cats of all the major feline viruses which include feline immunodeficiency virus and feline coronaviruses (11). Although multiple species of felids are susceptible to a FeLV infection, only domestic and wild cats of the *Felis* genus harbor endogenous FeLV (enFeLV) in their genome (7). Endogenous FeLV has been correlated to resistance against exFeLV infections and is also a precursor to the FeLV-B recombinant subgroup (12, 13).

Recombination events between exogenous and endogenous FeLV is one factor in the formation of subgroups of exogenous FeLV, which includes FeLV-A, -B, -C, -D, -E, and -T (3-5, 9, 14-16). The most epidemiologically relevant subgroup, FeLV-A, is an exclusively exogenous virus and is horizontally transmitted through social behaviors, via contact with blood, feces, and during suckling (13, 17, 18). The remaining have arisen via mutations, deletions, and recombination between enFeLV (or ERV-DC) and exogenous FeLV-A (exFeLV), in the envelope (*env*) gene (7, 9, 15, 17, 19, 20); FeLV-B is by far the most common recombinant variant described. Recombination in *env* allows for a change in cellular tropism and result in a different disease progression. Although these recombination events and mutations in *env* have typically been associated with increased pathogenicity, these FeLV variants are associated with decreased ability for horizontal transmission and require FeLV-A as a helper virus (9, 16, 17, 20-22). For example, horizontal transmission of FeLV-B has been documented in three different cases among many putative emergent infections (23, 24). Therefore, it is thought that with rare exception, each enFeLV-FeLV-A recombination event occurs *de novo* and is unique to each infected individual (9, 20, 21).

While the presence of FeLV-B has been well documented, little work has been done to understanding recombination events that predispose to FeLV-B emergence during natural infection. Defining the molecular underpinnings of this process provides a better understanding of the molecular basis for FeLV-B disease and provides important mechanistic details about endogenous-exogenous retroviral interactions (25). We therefore genotyped naturally emergent FeLV-B isolates from a closed privately-owned colony of domestic cat with epizootic FeLV (13) to investigate how FeLV-B infection arises and spreads within a population. We hypothesized four scenarios which could be tested by our analysis (**Figure 1A-D**): (A) No recombination events (i.e., no FeLV-B detected), (B) FeLV-B formation resulting exclusively via *de novo* recombination within individuals, (C) FeLV-A+B arising in a minority of individuals and subsequently transmitted horizontally, or (D) FeLV-B *de novo* emergence occurring simultaneously with horizontal transmission. We analyzed FeLV-A, FeLV-B, and enFeLV *env* sequences from natural FeLV-A and FeLV-B infection to assess phylogenetic relatedness and recombination breakpoints from 22 individuals to determine relationships among these agents.

**Figure 1.**
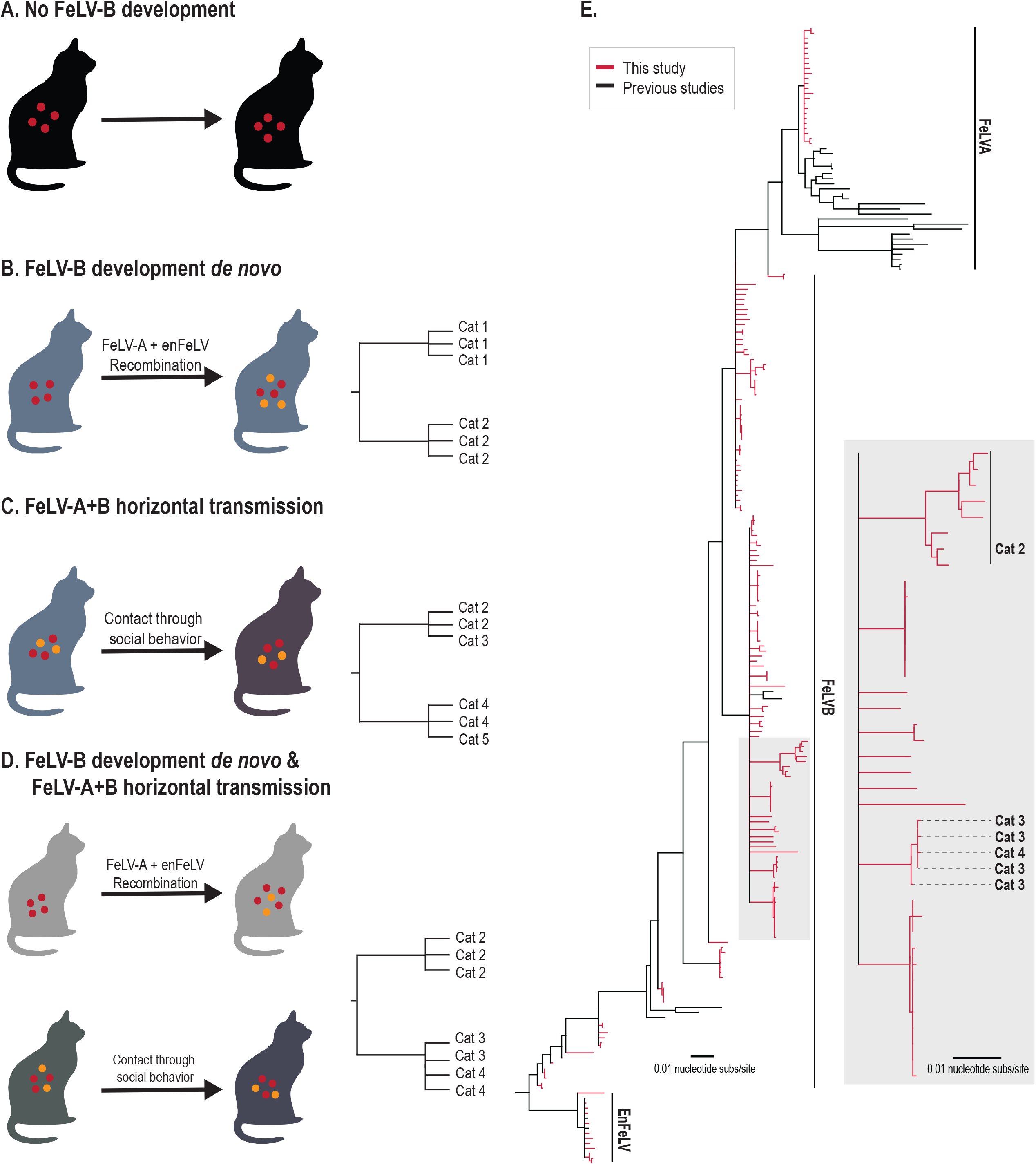
FeLV-B phylogeny reveals a high degree of genetic dissimilarity within the closed cat colony, and supports *de novo* recombination as a primary source of FeLV-B emergence. **A-D**. Four potential scenarios for FeLV-B emergence were considered which could be distinguished by phylogenetic analysis. **A**. No recombination. A cat with active FeLV-A infection (red circles) that does not develop into FeLV-B. Ten of 32 cats evaluated were in this category. **B**. A cat has an active FeLV-A infection (red circles) which recombines with enFeLV (yellow circles) to develop a FeLV-B (orange) infection. If all FeLV-B infections developed *de novo* and horizontal transmission was not possible or rare, genomic analysis would cluster viral sequences from the same individual together or show recombination events that are not identical between individuals. **C**. A cat horizontally transmits both FeLV-A (red circles) and FeLV-B (orange circles) to another cat via social contact. In this case, phylogenies would indicate branches with shared FeLV-B ancestry between infected individuals. **D**. FeLV-B infection (orange circles) emerges via both *de novo* recombination and horizontal transmission. Resulting phylogeny would demonstrate both independent FeLV-B branches related to each cat as well as mixed branches indicating FeLV-B infections shared between individuals. It is important to note that different FeLV-B recombination break points that occur within an individual could be shared across individuals and would generate a similar phylogenic structure as direct transmission of FeLV-B and could confound interpretations. **E**. Neighbor-joining phylogenetic tree illustrates the relationship between FeLV sequences (FeLV-A, B and enFeLV) recovered in this study (red) and sequences reported previously (black). Two primary clades and several minor clades were identified. Several FeLV-B clades are seen here, which are indicative of the differing recombination patterns between FeLV-A and enFeLV. This results in FeLV-B isolates that are more closely related to FeLV-A, isolates that are more ‘enFeLV-like’, or variants which are intermediate between these two identities. Our data suggests scenario D though the majority of FeLV-B variants arising *de novo*.

## Results

### Prevalence of FeLV within the colony

We previously reported that 32 cats from a colony with 65 individuals were diagnosed with FeLV antigenemia by a commercially available point-of-care test (49.2%; 13). Results were verified using a FeLV-A specific polymerase chain reaction (PCR). FeLV-B-specific PCR determined that 22 of 32 FeLV-A positive individuals had detectable FeLV-B infection (68.8%, FeLV-A+/B+). No individuals were FeLV-B positive without a concurrent FeLV-A infection (FeLV-A-/B+; 13).

### Genetic analyses of FeLV subgroups

#### FeLV-A and enFeLV sequences collected are nearly identical in all cats

One hundred and ninety-two FeLV sequences were analyzed from the *env* region of FeLV-A (22 sequences from 16 individuals), FeLV-B (156 sequences from 22 individuals) and enFeLV (14 sequences from 14 individuals) were sequenced from domestic cats (**Table 1**). Phylogenetic analysis of this dataset indicated that all FeLV-A sequences collected from the colony are highly similar and share 99-100% identity (**Figure 1E**, pairwise data not shown). As anticipated, enFeLV sequences, which are fixed in the domestic cat genome, were 98-100% similar (**Figure 1E**, pairwise data not shown). FeLV-B sequences were more variable, with 88-100% sequence identity (**Figure 1E**, pairwise data not shown). FeLV-B phylogeny documented emergent viruses in this colony form multiple clades. Specifically, FeLV-B variants could be grouped into clades that were ‘FeLV-A-like’, ‘enFeLV-like’ or intermediate (**Figure 1**).

**Table 1:**
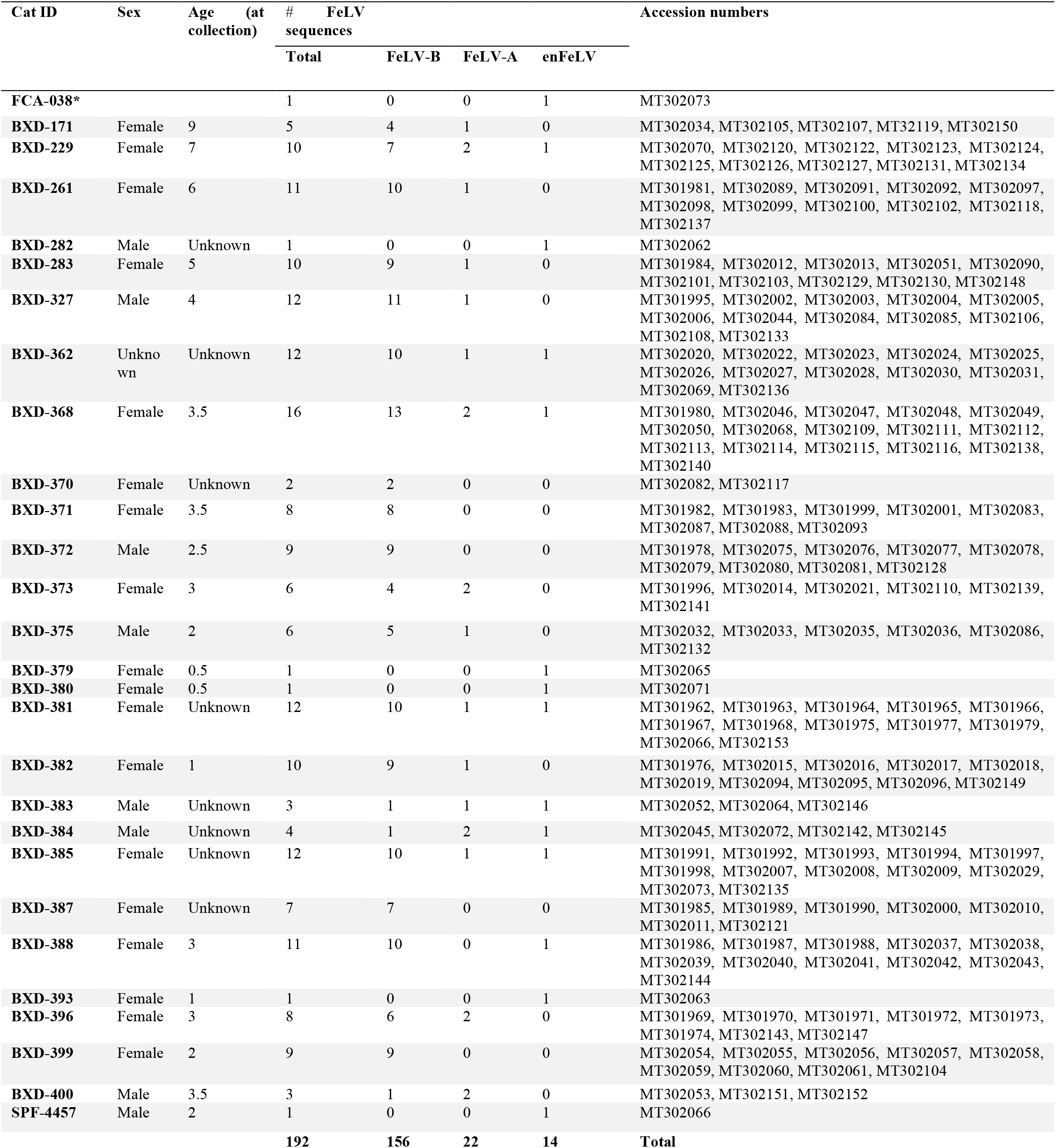
Sample information including CatID, demographics, FeLV group and accession number.

#### FeLV-B recombination sites were variable

Recombination Detection Program 4 (RDP4) analysis identified numerous recombination breakpoint sites. A large number of recombination events were identified. Due to the similarities in sequence identity, many recombination events among FeLV-A, endogenous FeLV, and FeLV-B contribute to the genetic diversity of FeLV-B. For the purpose of this study, we only highlighted those events which specifically were detected to have occurred between enFeLV and FeLV-A (resulting in FeLV-B variants). Our analyses detect recombination sites within p15E and the 3’ end of gp70 (**Figure 2**). The majority of sites were located in the 5’ end of gp70. We were unable to detect all potential 5’ recombination sites due to the location of our forward primer, and because of complexity of recombination patterns in this region. FeLV-A and enFeLV recombination sites are surprisingly numerous, resulting in a large number of FeLV-B variants. A tanglegram analyses of FeLV-A and FeLV-B variants from single hosts determined that variants are not unique to a single host (**Figure 3**). Of 18 individuals where both FeLV-A/B sequences were recovered, nine (50%) had multiple FeLV-B variants (**Figure 3**). While several FeLV-B sequences from this study cluster with those sequenced previously, we were able to identify multiple unique clades within this single colony (**Figure 1**).

**Figure 2.**
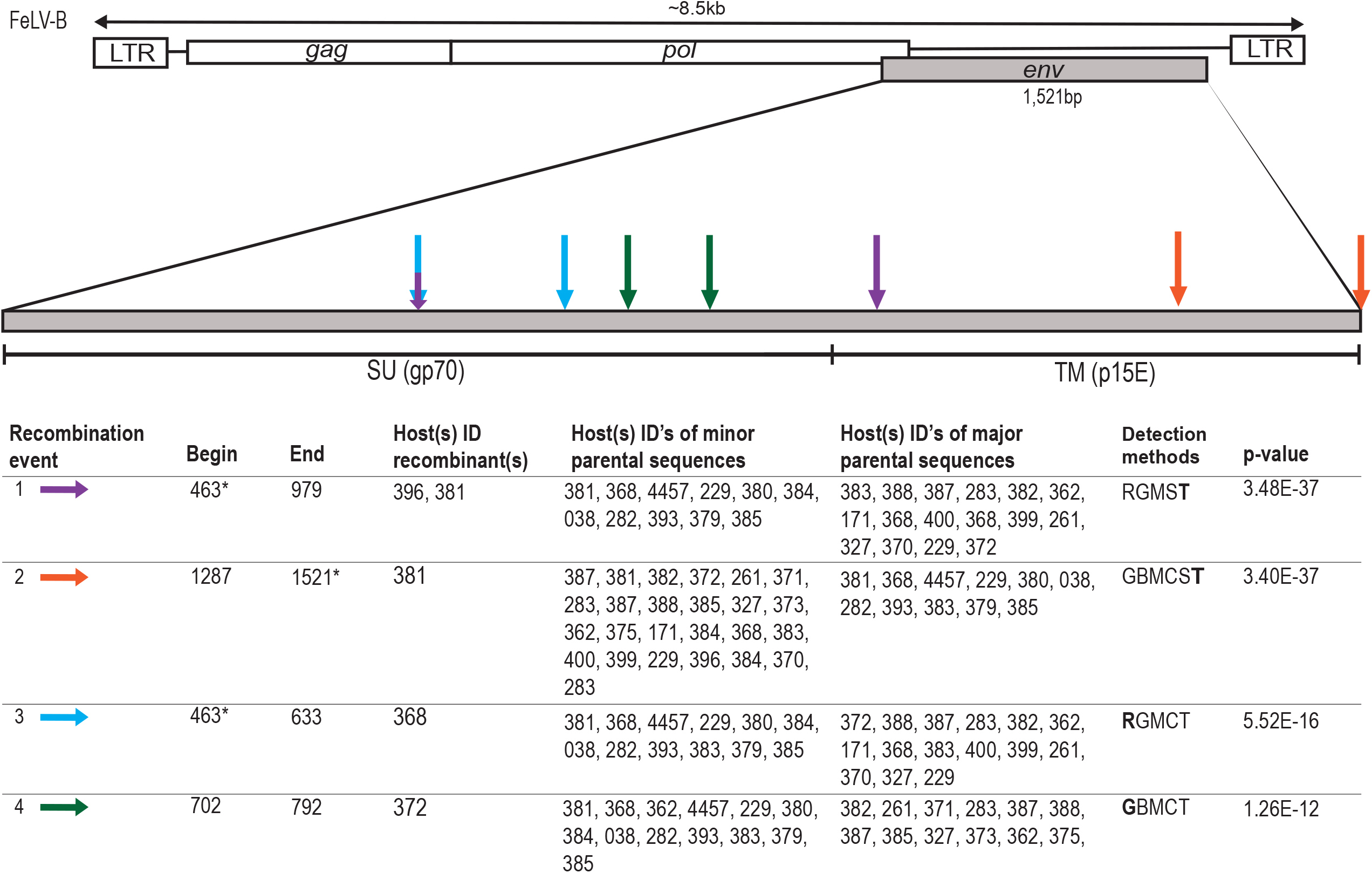
Multiple recombination sites in *env* identified among FeLV-A, enFeLV, and FeLV-B isolates. Schematic illustrates recombination events in FeLV-B inferred from parental FeLV-A and enFeLV sequences. The majority of sites were concentrated in the 5’ end of gp70. Recombination sites for each event are represented by arrows of different colors. Information below schematic shows recombination information including event, host ID harboring recombinant sequence, host ID harboring parental sequence, detection methods with best method (bold) and the corresponding highest e-value.

**Figure 3:**
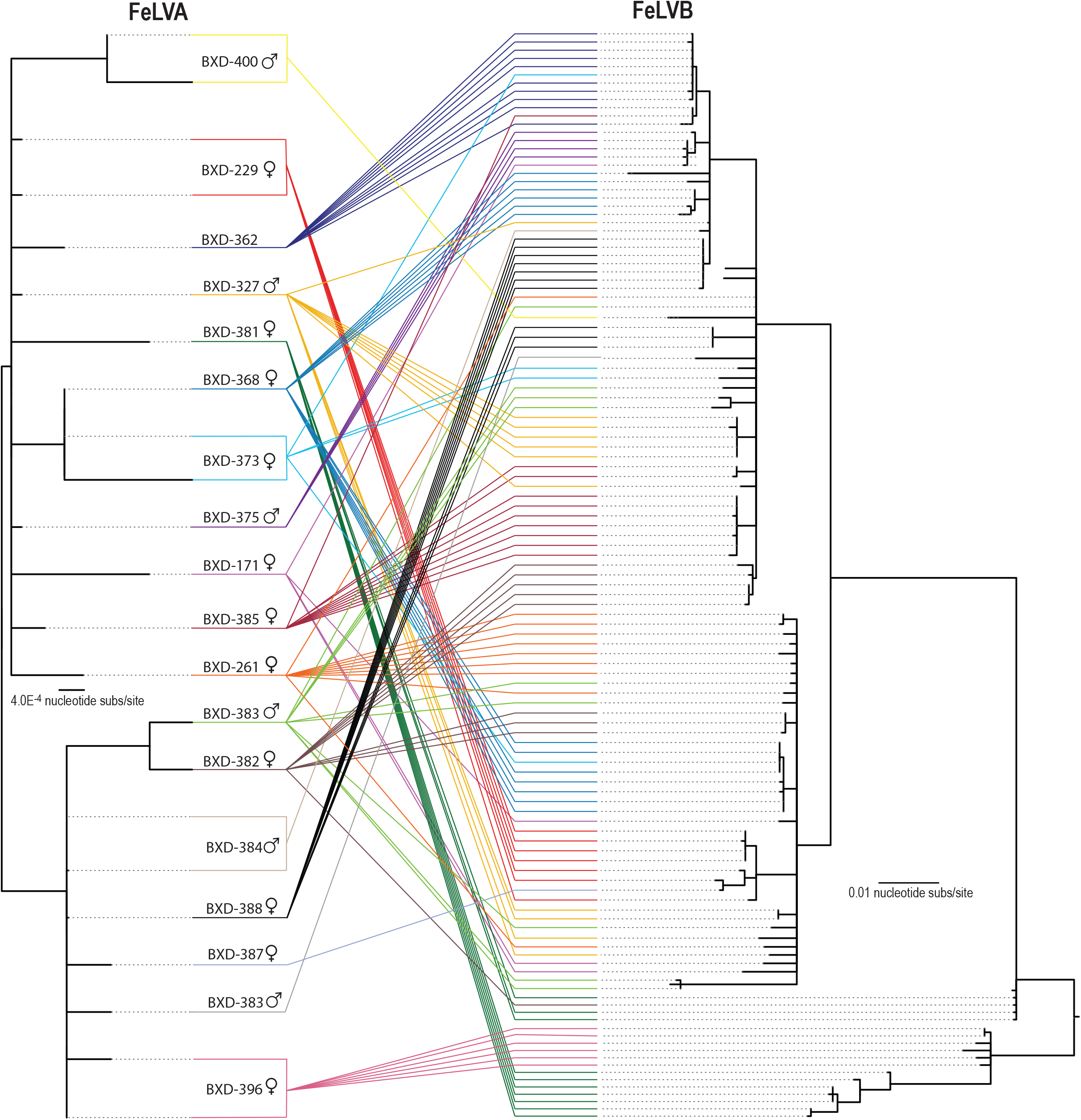
Comparison of FeLV-A and FeLV-B sequences from individual cats illustrates both de novo recombination and horizontal transmission. Neighbor joining phylogeny on the left illustrates FeLV-A *env* recovered from a single individual. Tanglegram illustrates relationship with FeLV-B *env* recovered from the same individual (right). Colored lines highlight *env* of two subtypes recovered from the same individual. This figure demonstrates that *env* variants from an individual are usually closely related; however, in half the cats with multiple variants detected, FeLV-B variants were distinct enough to group in separate clades.

## Discussion

This study of FeLV-B variants emerging during natural FeLV-A infection in a closed colony of domestic cats has important implications not only for pathogenesis of FeLV disease in cats, but in furthering our general understanding of endogenous-exogenous viral interactions (26). The most thoroughly characterized ERV-XRV interactions have been described in mice (mouse mammary tumor virus, murine leukemia virus), sheep (jaagsiekte sheep retrovirus), chickens (Rous sarcoma virus, avian sarcoma leukosis virus), koalas (koala retrovirus), and in cats with FeLV (27). Few studies have been performed in times of modern genomics analysis, particularly in infections occurring in a natural setting that have been well-characterized clinically and virologically, and thus our findings provide novel insights that lay the groundwork for future studies.

Although enFeLV is defective and incapable of forming infectious virions, open reading frames in several genomic segments, most notably *env*, result in full-length transcripts in certain tissues (19). FeLV-A recombination with enFeLV occurs during retroviral transcriptase-directed DNA synthesis following co-packaging of endogenous and exogenous viral genomes (28). Our findings of common but varied sites of recombination concentrated in the N-terminal portion of *env* support previous results (12, 15). These hotspots likely represent locations within *env* where template switching between FeLV-A and enFeLV is favorable. The resultant recombinant FeLV-B are more virulent; however these variants are replication-defective and require an FeLV-A helper virus for successful replication. The variation in recombination sites often result in changes in pathology or infectivity of FeLV-B associated with ability to utilize new receptors. For example, certain FeLV-B variable region A (VRA) sequences within the receptor binding domain facilitate binding to both Pit 1 and Pit 2 receptors (29). Conversely, some variants may also result in a non-functional recombinant. The FeLV surface glycoprotein (SU gp70) is often under strong selective pressure from host immune response as it is responsible for binding to host cell receptors and a common target of neutralizing antibodies (26, 30, 31). This selection has resulted in extensive genetic variation in FeLV SU coding regions (26), which is consistent with our findings of multiple recombination sites at a variety of locations with the SU gp70 region. The frequency and variation of recombination events in this region suggest selective pressures have resulted in mechanisms for greater viral variation in this region. One event was identified in the transmembrane (TMp15E) region and another that spans much of the SU and the 5’ region of the transmembrane. The functional changes associated with these recombination sites is unknown but previous work using chimeric viruses has established that differences in amount of enFeLV in SU gp70 alters viral replication efficiency, cell tropism, and tendency to induce aggregation of lymphoid cells in tissue culture (12). This study investigates these variant patterns in an individual cats living in a closed cat colony, validating previous *in vitro* results (12), and illustrating the extent that within and between individual recombination between FeLV-A and enFeLV occurs, resulting in high diversity in key coding regions of FeLV-B. It is notable that the recombination patterns in the SU gp70 region correlate with those predicted by in vitro experiments nearly 30 years ago by Pandey et al (12).

Understanding the emergence of FeLV-B has important implications to feline health as emergence of this recombinant virus is associated with poorer health outcomes and higher viral loads (13, 14, 22). For example, should FeLV-B be readily transmitted amongst cats, it would indicate that strain specific testing may be important prior to housing infected cats together. Alternatively, if FeLV-B only forms within individual cats from recombination between enFeLV and FeLV-A, it would indicate this is a frequent occurrence that could increase variation of FeLV should recombination occur at different locations. Our findings suggest that the FeLV-B infections arise *de novo* within individual cats, as no cats were singularly infected with FeLV-B and the majority of FeLV-B infections from a single cat clustered together phylogenetically. A tanglegram comparing FeLV-A with multiple clones of FeLV-B from the same cat further supports variation in FeLV-B is a primarily a function of FeLV-A variation. However, we identified 9 cats with more than one distinct FeLV-B variant, indicating that FeLV-B may at times be co-packaged and transmitted with FeLV-A, and/or that more than one recombination event can emerge in an individual cat. While variation in recombination sites complicates phylogenic analysis of FeLV-B and thus assessment of cat to cat transmission of FeLV-B is inconclusive, we have documented some shared variants between cats suggesting that horizontal transmission is possible. Further, we have previously reported concurrent FeLV-A/FeLV-B cross-species transmissions in Florida panthers lacking enFeLV, supporting the potential for horizontal FeLV-A/FeLV-B co-transmission events (23). The large number of unique FeLV-B sequences detected strongly supports that FeLV-B predominately arises *de novo*, and variation in FeLV-B is largely driven by FeLV-A recombination site promiscuity.

Finally, while the majority of cats in this colony with progressive FeLV-A infection were FeLV-B positive, one third of the cats were FeLV-B negative, consistent with other reports that FeLV-B does not emerge in every case of FeLV-A infection (14). Examining cellular mechanisms that relate to resistance to FeLV-B emergence, and concurrently better prognosis, is an area for future studies to understand factors restricting ERV-XRV recombination.

## Materials and Methods

### Animals and sampling

Blood samples were collected in EDTA from 65 individual cats (leopard cat × Domestic cat cross) in a privately-owned breeding colony at a single time point and shipped overnight on ice. The colony consists of domestic cat-Asian leopard cat (*Prionailurus bengalensis*) hybrids that have been backcrossed with domestic cats for several generations. At time of blood collection, animals ranged in age from 8 weeks to 9 years with 30 males, 32 females, and 3 animals of unknown sex. All procedures and handling of samples were done by a licensed veterinarian (13).

### Sample processing

DNA was isolated from Peripheral blood mononuclear cells (PBMCs) using a DNeasy blood and tissue extraction kit (Qiagen, Inc, Valencia, CA). DNA concentration was determined using a NanoDrop-1000 Spectrophometer (Thermo Fisher Scientific, Waltham, MA, USA). Blood sampled were stored at -80°C and working samples stored at -20°.

### FeLV Status

During a previous study commercially available enzyme-linked immunosorbent assay (ELISA) SNAP tests (IDEXX Laboratories, Westbrook, ME, USA) were used to determine FeLV status using plasma and serum samples (13). FeLV-A and FeLV-B status was established using a FeLV specific PCR (described below).

### Amplification of FeLV-A, -B and enFeLV sequences

A PCR protocol was developed to target the *env* gene and LTR region of FeLV-B by developing primers that would specifically only bind to FeLV-B recombinant sequences. To accomplish this the following primers were used: forward primer (5’-CAGATCAGGAACCATTCCCAGG-3’) specific for enFeLV in a known recombination site in the *env* gene, and reverse primer (5-CCTCTATCTTCCTTGTATCTCATGG-3’) specific for exFeLV in the LTR region. Given the LTR region of FeLV-B is derived from exFeLV, we used forward primer (5’-ACCCAAGCTAATGCCACCTC-3’) specific for FeLV-A in the *env* gene and reverse primer (5’-CCTCTAACTTCCTTGTATCTCATGG-3’) specific for FeLV-A in the LTR region. Specific enFeLV primers were designed to target an already known enFeLV recombination site not present in exFeLV variants, forward primer (5’-CTGACAGACGCCTTCCCTAC-3’) specific for enFeLV in the *env* gene and reverse primer (5’-CTAGGCTCATCTCTAGGGTCTATC-3’. These primers were using in a PCR reaction with the following specifications; each reaction contained 1µL of 10µM concentration of forward and reverse primer, 10µL of Kapa HiFi mastermix (Kapa Biosystems), 7µL of water, and 2µL of DNA template in a 20µL reaction. Thermal cycling conditions were 95°C for 3 min., followed by 30 cycles of 95°C for 30 sec., 58°C for 30 sec., 72°C for 2 min., ending with 72°C 3 min. Each set of PCR reactions included negative controls to assure positive reactions were FeLV-B specific, namely specific pathogen free (SPF) cat DNA containing enFeLV sequences (to ensure that enFeLV was not the amplification product), and a no template control (DNA free water). A confirmed FeLV-B sample from the colony was included as a positive control. Positive PCR samples were run on a 0.7% agarose gel at 80V, 400mA for 30-45 minutes. Bands of interest were excised and DNA was extracted using MEGAquick-spin™ plus fragment DNA purification kit (iNtRON Biotechnology, South Korea). Purified DNA was ligated into a Pjet1.2 plasmid and plasmids were then isolated using DNA-spin™ plasmid DNA purification kit (iNtRON Biotechnology, South Korea) which were then sequenced by a commercial laboratory using Sanger protocols (Macrogen Inc, Rockville, MD). One to thirteen clones from each host were sent, depending upon sample availability and number of colonies obtained.

### Sequence analysis

Sequences were trimmed and aligned in Geneious (Biomatters Inc, Newark, NJ) with representative FeLV subtypes downloaded from Genbank. Initial analysis confirmed grouping with FeLV-B/FeLV-B-like, FeLV-A, or enFeLV-like sequences. Datasets were compiled of those sequences recovered in this study and representatives available in Genbank from the three FeLV groups using the software Molecular Evolutionary Genetic Analysis (MEGA) (32) and aligned using MUSCLE (33). This dataset was used to construct a neighbor-joining phylogenetic tree using jukes-cantor model and 1,000 bootstrap replicates in MEGA (32). Completed trees were then uploaded to iTOL (interactive tree of life) (34), which allowed for the trees to be further modified. This comparison was used to inform the spread of FeLV-B infection vs FeLV-A infection within a single individual as well as between infected individuals in the study group.

### Recombination analysis

Recombination Detection Program 4 (RDP4) was used to detect recombination sites within FeLV-B (35). RDP (Martin and Rybicki, 2000), GENECONV (Padidam et al., 1999), Bootscan (Martin et al., 2005), Maxchi (Smith, 1992), Chimera (Posada and Crandall, 2001), Siscan (Gibbs et al., 2000), and 3Seq (Boni et al., 2007). Recombination events were only accepted when they were detected by three or more of these methods with associated p-values <10^3^ with strong phylogenetic support. Sequences also underwent analysis by Sequence Demarcation Tool version 1.2 (SDTv1.2) (36). Tanglegram consists of neighbor-joining phylogenetic trees of the FeLV-A and FeLV-B datasets from those cats where sequencing of both variants were recovered, constructed using the parameters described above.

## Acknowledgements

This work was supported by NSF-EID award 1413925 (VandeWoude) and by the Office of the Director, National Institutes Of Health of the National Institutes of Health under Award Number F30OD023386 (Chiu). The content is solely the responsibility of the authors and does not necessarily represent the official views of the National Institutes of Health. The funders had no role in study design, data collection and interpretation, or the decision to submit the work for publication.

